# Installation of Dominant-Negative Mutations in *FAS* and *TGFβR2* via Base Editing in Primary T Cells

**DOI:** 10.1101/2025.09.11.675648

**Authors:** Bryce J. Wick, Mitchell G. Kluesner, Nicholas J. Slipek, Joseph G. Skeate, Ethan M. Niemeyer, Beau R. Webber, Branden S. Moriarity

## Abstract

Adoptive cell transfer (ACT) of engineered T cells is effective against B-cell malignancies but has faltered against solid tumors due to the immunosuppressive tumor microenvironment (TME). FASL and TGFβ are key mediators of T cell dysfunction in the TME and overexpressing dominant negative (dn) forms of their receptors in T cells increases anti-tumor efficacy in solid tumor models. However, an approach which directly targets the endogenous genes would be more amenable to multiplex editing and reduce competition with WT alleles. Here, we employ base editing (BE) in primary human T cells to install naturally occurring dominant negative *FAS* and *TGFβR2* mutations. *In vitro* survival and proliferation assays demonstrate that BE T cells are resistant to pro-apoptotic and anti-proliferative effects of FAS and TGFβ signaling. CAR-T cells with BE-installed dn TGFβR2 or dn FAS exhibit improvements in cytotoxicity, while dn TGFβR2 CAR T demonstrate increased persistence and reduced expression of phenotypic markers of exhaustion compared to controls. Moreover, BE-engineered dn CAR T outperform lentiviral-engineered cDNA over expression counterparts in several functional assays. Considering the efficiency of BE and its amenability for multiplex editing, our novel approach lends itself to engineering strategies necessary to overcome T cell dysfunction in solid tumors.

## Introduction

Adoptive cell transfer (ACT) using chimeric antigen receptor (CAR) T cells has demonstrated promising efficacy when deployed against CD19+ and BCMA+ hematologic malignancies ^1–3^. However, the same cannot be said for solid tumors, where CAR-T therapy has been comparatively ineffective ^4–6^. As we continue to uncover the underlying mechanisms of CAR-T failure in solid tumors, the immunosuppressive tumor microenvironment (TME) has emerged as a driving factor ^7,8^. Two critical ways in which the TME suppresses effector cell function are through the transforming growth factor beta (TGFβ) and FAS pathways ^9–12^. Many cancers are known to secrete high levels of soluble TGFβ, which inhibits T cell proliferation and cytokine production in the TME ^13,14^. FAS ligand (*FASL)* is expressed on many cancers, which binds to its cognate receptor FAS expressed by T cells resulting in apoptosis and diminished efficacy ^15^. Efforts to generate CAR-T cells resistant to FAS and TGFβ signaling have armored them against the immunosuppressive TME and increased their persistence, resilience, and efficacy ^15–17^. The primary method that has been deployed to abrogate FAS and TGFβ signaling is through the overexpression of dominant-negative (dn) receptors ^15–17^. These dn receptors have been altered via truncation or mutation of the intracellular domains, such that even though the receptor still binds cognate ligands, it is not capable of transducing downstream signals. Thus, dn receptors can not only eliminate downstream signaling, but also sequester ligands that could suppress other immune effector cells.

Currently, the most common method to introduce dn receptors in CAR-T cells is to express them via the introduction of a cDNA using viral vectors. This method has two major drawbacks. One is that the expression and function of endogenous receptors is not altered, allowing signaling to continue, albeit at a lower level ^18,19^. Moreover, the inclusion of additional genetic cargo increases vector size, reducing viral titers, expression levels of CAR constructs, and in many cases is not possible due to the packaging limits of viral vectors ^20,21^. A method that bypasses viral vectors by directly modifying the endogenous gene, supports higher-order multiplex editing, and eliminates residual signaling while preserving CAR expression would enhance the efficacy and capability of CAR T products.

Base editors can introduce single nucleotide changes with high efficiency and product purity and without double-stranded DNA breaks (DSBs), obviating the requirement for a DNA donor molecule ^22^. Here, we deploy base editing (BE) to introduce naturally occurring, dominant-negative point mutations in *FAS* and TGFβ receptor 2 (*TGFβR2*) to disrupt their respective signaling pathways. Our approach efficiently converts endogenous FAS and TGFβR2 receptors to dn forms *in situ*, without the need for incorporation of additional cDNAs or induction of DSBs. This approach has not been demonstrated previously, as base editors have not been used to introduce dominant-negative mutations. This strategy lends itself to clinically focused CAR-T manufacturing, especially in the context of multiplex engineering strategies that are becoming increasingly recognized as important to overcome the immunosuppressive TME in solid tumors.

## Results

### Base editors efficiently install dn mutations at endogenous loci in primary human T cells

To identify candidate sites for installation of dn mutations, we assessed naturally occurring point mutations known to disrupt signaling. For *FAS*, we surveyed point mutations linked to autoimmune and lymphoproliferative syndromes due to their reported abrogation of FAS signaling^23–27^. For *TGFβR2*, we surveyed function abrogating point mutations associated with either Loeys–Dietz syndrome or breast cancer ^28–32^. To this end, we designed 12 gRNAs, 6 per gene, including a gRNA targeting a splice acceptor sequence as a traditional KO control ^23,33^ **(Figure 1A, B)**. These mutations were primarily in the intracellular region of the receptors, with the exception of FAS T28A; and all mimic natural dn mutations affecting signal transduction without impacting ligand binding **(Figure 1A)**. In instances where the exact naturally occurring amino acid change could not be installed with adenosine base editing (A->G), we introduced a substitution of the target amino acid predicted to recapitulate disruption of endogenous receptor signaling. To test the feasibility of generating these edits, we electroporated primary human T cells with *FAS* and *TGFβR2* targeting gRNAs and adenine base editor (ABE8e) ^24^ mRNA and assessed BE efficiency via Sanger sequencing 7 days post electroporation. We used ABE8e as it is a highly active and precise BE with low off target activity and high product purity ^24,34^. Most gRNAs achieved efficient targeted A->G base conversion, with three gRNAs for *FAS* and four gRNAs for *TGFβR2* demonstrating >50% base conversion of the target nucleotide **(Figure 1C, Supplemental Figures S1 and S2)**. These results demonstrate that dn mutations informed by naturally occurring mutations can be efficiently installed in primary human T cells with BE technology.

**Figure 1.**
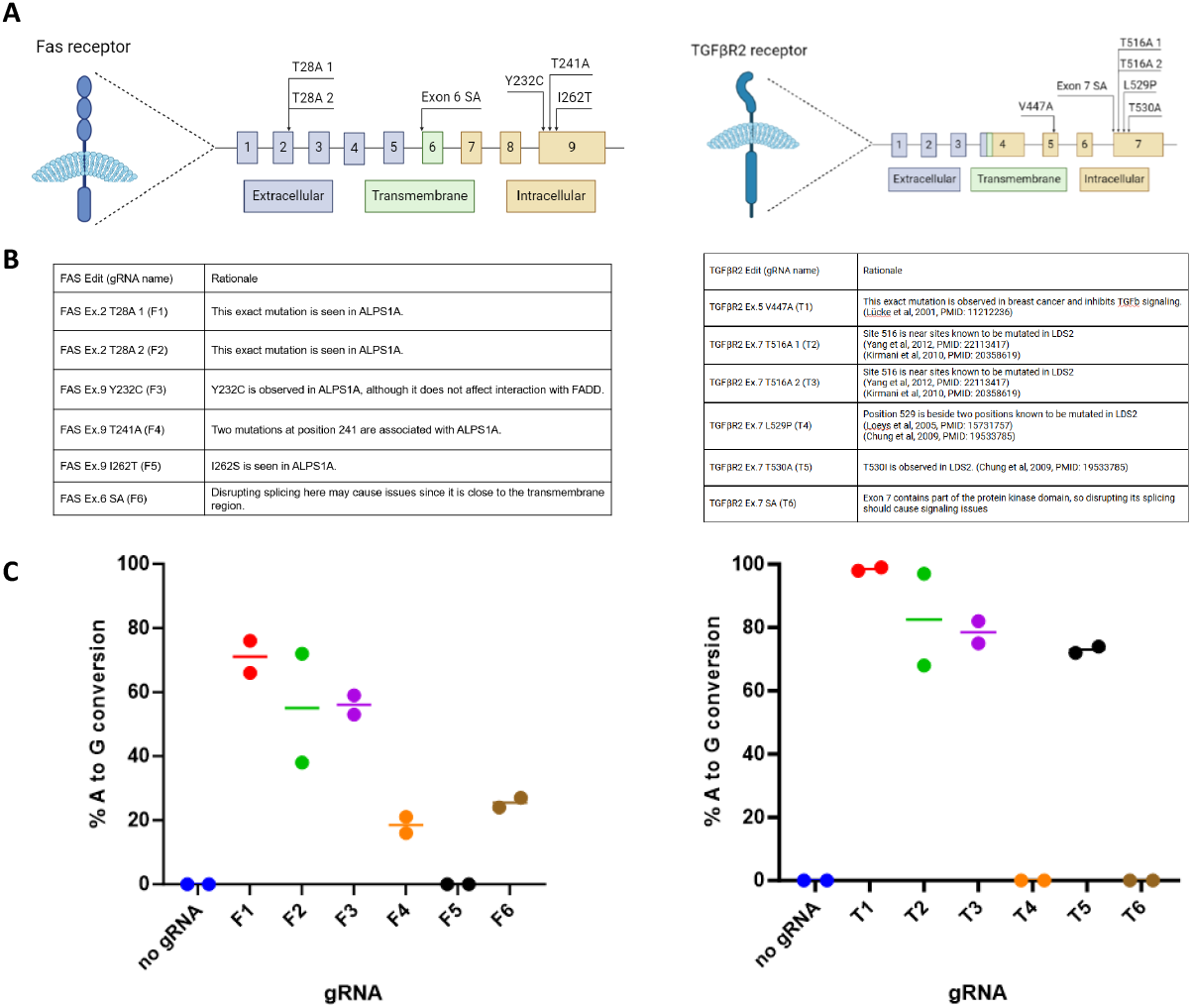
Base editor technology can efficiently install dn mutations in primary human T cells. (A) Gene diagrams displaying location of each gRNA target site. (B) Tables describing rationale behind each gRNA. Guide RNAs are designed to either create specific mutations seen in naturally-occurring disease associated with receptor dysfunction or to target splice sites. (C) Base editing activity of each gRNA, FAS on the left and TGFβR2 on the right. Guide RNAs were electroporated into T cells with ABE8e mRNA and base editing activity was assessed seven days later via sanger sequencing. (n = 2 donors).

### T cells harboring dn receptor mutations installed by BE are resistant to suppression by FASL and TGFβ

To confirm that BE T cells are indeed resistant to FASL and TGFβ signaling, we conducted *in vitro* functional assays for both dnFAS and dnTGFβR2. For the dnFAS assay, engineered T cells were exposed to FASL and a His-Tag antibody for 24 hours **(Figure 2A)**. The His-Tag antibody allows FASL to naturally trimerize for proper signaling and induction of apoptosis in FAS expressing cells ^25^. Cell viability was measured after 24 hours, and of the three gRNAs with efficient base conversion **(Supplemental Figures S3A and S4)**, only FAS Y232C generated resistance to FASL mediated apoptosis. Specifically, we noted significantly reduced T cell death by ~80% compared to unedited controls **(Figure 2B)**. To see if the reduction in cell death was in fact due to reduced FAS signaling, we analyzed a downstream signaling molecule of the FAS pathway, namely cleaved Caspase 3 (cC3). After confirming high levels of base conversion **(Supplemental Figures S3B and S5)**, intracellular staining (ICS) and analysis by flow cytometry showed that FAS Y232C led to a ~75% reduction in cC3 after 24-hour exposure to FASL relative to unedited controls **(Figure 2C, Supplemental Figure S3C)**.

**Figure 2.**
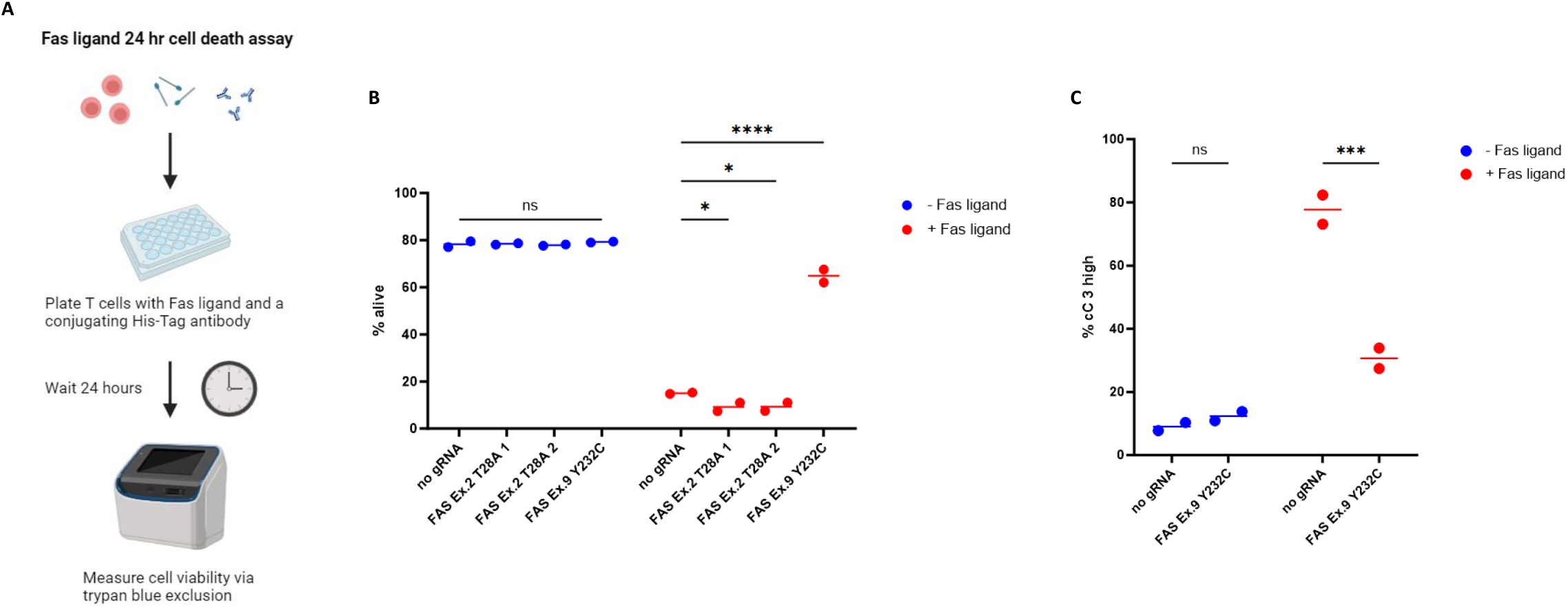
FAS Y232C reduces FAS signaling activity and promotes resistance to FASL-mediated apoptosis. (A) Diagram of Fas ligand-based killing assay. (B) Percent live cells remaining after Fas ligand-based killing assay. (C) ICS of downstream protein target. T cells were either exposed to Fas ligand and a conjugating His-Tag antibody for 24 hours or just the His-Tag antibody and then ICS was performed for cleaved Caspase 3. (n = 2 donors). ns = not significant, *p<0.05, **p<0.01, ***p<0.001, ****p<0.0001, (B) Two-way ANOVA with Dunnett’s multiple comparisons test against no gRNA control. (C) Two-way ANOVA with Uncorrected Fisher’s LSD.

To functionally validate candidate dnTGFβR2 mutations, we designed a serial stimulation experiment to assess TGFβ-mediated inhibition of T cell proliferation in wild-type and BE T cells. Briefly, T cells were repeatedly stimulated with anti-CD3 and anti-CD28 antibodies in TGFβ-containing media and cell proliferation was measured over the course of repeated stimulations **(Figure 3A)**. We used G-Rex plates for minimal T cell disturbance and rested T cells 11 days without anti-CD3/CD28 stimulation before restimulation rounds. Only one TGFβR2 gRNA led to significantly improved proliferation relative to controls, TGFβR2 V447A. After three rounds of stimulation, the V447A mutation demonstrated a significant 2.8-fold increase in total cell numbers compared to controls with a 2-fold increase in overall viability **(Figure 3B, Supplemental Figure 6A)**. To confirm that this phenotype is due to disrupted TGFβ signaling, we assessed downstream signaling molecules in the TGFβ pathway, namely SMAD2/3. After confirming high levels of base conversion **(Supplemental Figures S3B and S5)**, ICS demonstrated that TGFβR2 V447A led to a 92% reduction in SMAD2/3 expression following exposure to TGFβ relative to unedited controls **(Figure 3C, Supplemental Figure S6B)**. These data demonstrate that BE installed Y232C and V447A mutations functionally disrupt FAS and TGFβ signaling, respectively.

**Figure 3.**
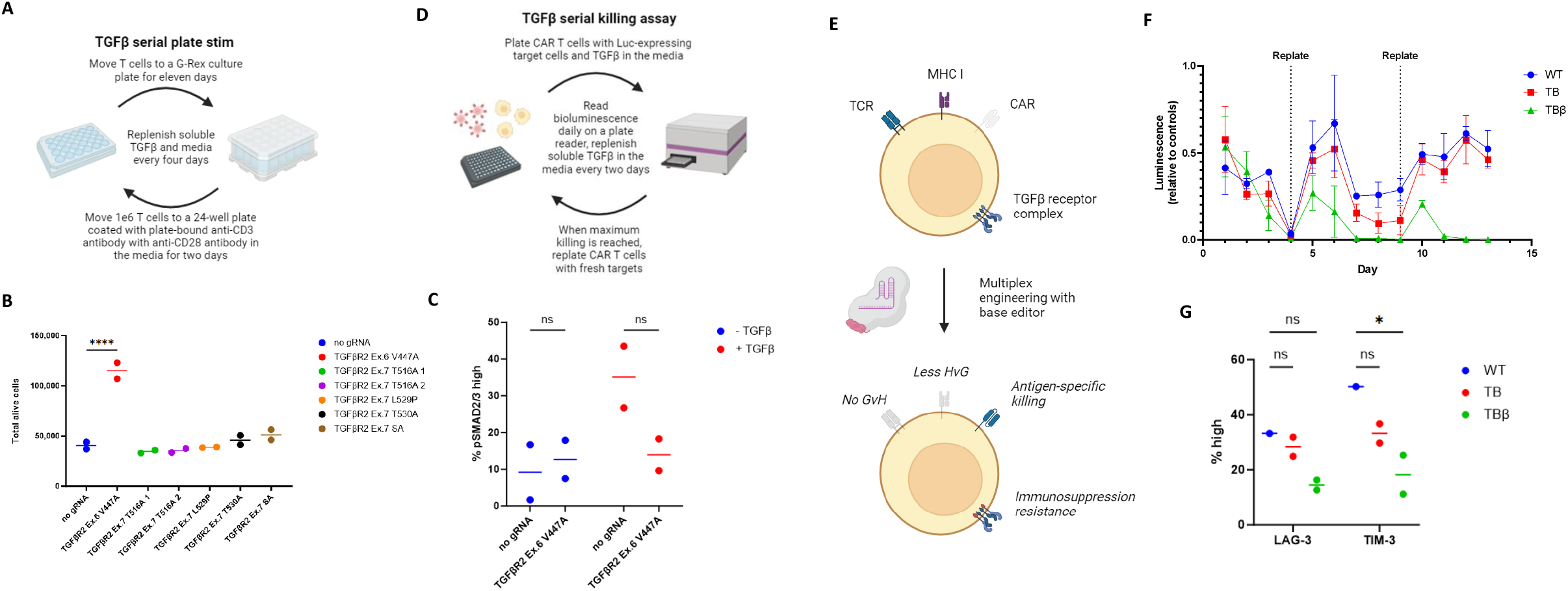
TGFβR2 V447A reduces TGFβ signaling activity and enhances CAR-T function in the presence of TGFβ. (A) Diagram of TGFβ serial plate stim assay. (B) Diagram of TGFβ serial killing assay. (C) Total live cells four days after the third stimulation in the serial plate stim experiment. (D) ICS of downstream protein targets. T cells were either exposed to TGFβ for 30 minutes or media alone and then ICS was performed for SMAD2/3. (E) Diagram explaining edits done to T cells and rationale behind each one for the TGFβ serial killing assay. (F) Serial killing assay results. Killing was measured via bioluminescence on a plate reader after D-Luciferin was added to the media. (G) Exhaustion panel done before the second replating during the serial killing assay. WT = wild type (no knock-outs). TB = TRAC KO and B2M KO. TBβ = TRAC KO, B2M KO, and TGFβR2 V447A. (n = 2 donors). ns = not significant, *p<0.05, **p<0.01, ***p<0.001, ****p<0.0001, (B) One-way ANOVA with Dunnett’s multiple comparisons test against no gRNA control. (C, E bottom) Two-way ANOVA with Uncorrected Fisher’s LSD.

### Dominant-negative TGFβR2 CAR-T cells demonstrate enhanced cytolytic function in the presence of TGFβ

To assess the functional impact of the V447A dn TGFβR2 mutation in a clinically relevant setting, we conducted a CAR-T cell serial killing assay in the presence of soluble TGFβ. Briefly, anti-CD19 CAR-T cells were co-cultured with Luciferase-expressing Raji target cells (CD19^+^) in the presence of TGFβ **(Figure 3D)**. Furthermore, to verify that dn receptor mutations are compatible with multiplex base editing, we also included previously validated BE gRNAs for KO of the immunologically relevant beta-2-microglobulin (B2M) and TCR (CD3e) in the CAR-T cells, **(Supplemental Figures S7A and S8)**. B2M KO may reduce host-vs-graft activity, prolonging the survival and activity of CAR-T therapy, while knocking out CD3e eliminates surface expression of the endogenous TCR in CAR-T cells, reducing potential off-target activity and graft-vs-host disease **(Figure 3E)** ^26^. In line with our previous data demonstrating efficient multiplex BE of T cells ^23^, including the extra gRNAs targeting CD3e and B2M in the CAR T engineering process did not significantly reduce CAR integration or TGFβR2 base editing rates, demonstrating the TGFβR2 V447A mutation can be engineered in a multiplex setting **(Supplemental Figures S7B, S7C, and S9)**. Installation of TGFβR2 V447A in CAR-T cells led to extended cytotoxic activity relative to controls in the serial killing assay **(Figure 3F)**. TGFβR2 V447A CAR-T cells not only killed target Raji cells faster and more effectively over rounds of serial killing, but also killed faster as the assay progressed to later rounds, demonstrating a clear advantage over control CAR-T cells in the presence of TGFβ. To determine if the cytolytic advantage exhibited by TGFβR2 V447A CAR-T cells was consistent with a less exhausted phenotype, we assayed a flow-based exhaustion marker panel on CAR-T before the last round of killing. Notably, TGFβR2 V447A CAR-T cells showed lower expression of both LAG-3 and TIM-3 relative to controls **(Figure 3G, Supplemental Figures S7D and S10)**. These data confirm that dn TGFβR2 mutations introduced at the endogenous locus using BE provide functional resistance to TGFβ suppression, representing a novel strategy to improve the resilience of CAR-T cells in an immunosuppressive solid tumor setting.

### BE-engineered dnFAS and dnTGFβR2 CAR T cells outperform lentiviral cDNA-engineered counterparts

As we are introducing a new engineering approach to install dn mutations, it is important to compare our new method to viral delivery of dn cDNAs, the most commonly used method of introducing dn receptors ^15–17,35,36^. To this end, we engineered dnFAS and dnTGFβR2 Mesothelin-CAR T cells from primary human T cells using a combination lentiviral delivery plus electroporation with ABE8e mRNA and gRNAs, as previously described ^23^, or through transduction using lentivirus containing a Mesothelin CAR and either dnFAS or dnTGFβR2 receptor cDNA^37,38^.

While both methods were able to introduce dnFAS and dnTGFβR2 with high efficiency as well as high CAR expression **(Supplemental Figures S11 and S12)**, the BE approach yielded higher dn receptor expression, nearing 100% while the lentiviral cDNA method reached 80% to 90% **(Supplemental Figure S12A)**. In a recombinant Fas ligand killing assay, the BE-engineered dnFAS CAR T cells demonstrated improved resistance to Fas ligand-mediated killing over lentiviral cDNA-engineered counterparts, reaching equal viability to non-FasL treated controls while the lentiviral cDNA-engineered cells did not reach the same level as their untreated controls **(Supplemental Figure S13A)**. Similarly, in a TGFβ serial plate restimulation experiment, the BE-engineered cells reached similar fold expansion levels to non-TGFβ treated controls while the lentiviral cDNA-engineered cells did not reach the same level as their untreated controls **(Supplemental Figure S13B)**. In killing assays against Mesothelin-expressing OVCAR-8 target cells, both BE and lentiviral cDNA methods to introduce dnTGFβR2 demonstrated improved killing in the presence of soluble TGFβ, although the BE-engineered CAR T cells outperformed the lentiviral cDNA-engineered cells. **(Supplemental Figure S13C)**. Finally, when dnFAS CAR T were subjected to a similar killing assay, BE-engineered cells once again outperformed their lentiviral cDNA-engineered counterparts **(Supplemental Figure S13D)**. Overall, this data demonstrates that when compared to lentiviral based cDNA over expression engineering, BE-mediated engineering methods result in CAR-T cells with higher resistance to inhibition.

## Discussion

Previously, we used BE for multiplex knockout to generate *PDCD1, B2M, and TRAC* deficient, off-the-shelf CAR-T cells with improved killing capability, increased expansion, and reduced translocations ^23^. Here, we demonstrate another exciting, novel application of BE to enhance cancer immunotherapy. Introducing dn mutations with BE has not been previously reported, and the applications of such an approach are wide-ranging, many of which are relevant well beyond the scope of CAR-T engineering. To design BE gRNAs that install dn receptor mutations, we surveyed naturally occurring mutations associated with disease as inspiration. However, considering FAS T28A is seen in ALPS1A, but in our experiments it did not yield any protection against FASL-induced apoptosis, a more reliable option for future dn mutation screening may be needed. Previously, we developed a bioinformatic tool to identify candidate BE gRNAs for splice site disruption ^33^. Similar programs also exist to aid in gRNA design for the introduction of premature stop codons with BE^17^. Considering the far-reaching potential dn mutations offer to a wide range of applications, a program for the design of gRNAs to install dn mutations would be immensely useful. For example, a recent screen done in primary human T cells with base editors mapped many mutations that tuned T cell function, but was limited by not including dn mutations ^39^. A tool to design dn mutation-installing gRNAs would allow for large scale dn mutation screens, accelerating novel mutation discovery.

Introducing dn mutations in endogenous receptors through BE offers key advantages over the current standard method for expressing dn receptors using cDNA ^15–17,35,36^. By using gRNAs to target endogenous genes instead of inserting whole cDNAs, multiplexing edits with this approach is significantly more feasible than with viral vectors.

Another advantage is that with cDNA expression, the endogenous receptors are still functional and reduce the overall efficacy of the dn receptor approach ^17,18^. Dominant-negative mutations also offer advantages over KOs, particularly in the context of incomplete KOs. This would be true for both single heterozygous cells where the dn receptors can sequester the functional receptors, and for a bulk population where dn cells can sequester ligands away from WT cells. Furthermore, a single dn point mutation may have less side-effects than a splice site mutation, which may cause bias for other isoforms of the target gene, leading to unintended consequences ^40^.

It should be mentioned that treating patients with CAR-T products engineered to harbor a mutation known to be associated with disease may carry enhanced risks. Therapeutic interventions using T cells, or any immune cell, engineered to be more active carry the risk of causing unwanted effects, such as hyperproliferation of the cellular therapy, or excessive cytokine production and subsequent toxicities. Therefore, extensive studies to ensure BE T cells do not overproliferate or secrete dangerous levels of cytokines would be needed during IND enabling drug studies. However, these kinds of concerns are not specific to dn mutations, as CAR-T engineered with randomly integrating vectors, KOs, or overexpression of various receptors or proteins carry similar concerns ^16^. Fortunately, the cancer immunotherapy field has already developed multiple avenues to combat these concerns, with many of them applicable to our dn mutation approach^18^. With this in mind, targeted *in situ* BE modification may offer a better safety profile than randomly inserting viral vectors or traditional Cas9 nuclease engineering. With no cDNA, site-specific base conversion, and no DSBs, *in situ* BE modification may lead to less direct genotoxicity over other current genetic editing technologies leveraged to enhance immune cell function.

In summary, our work describes an exciting new application of BE technology to enhance immune cell persistence and function. Our method is also multiplexable with other function-enhancing edits, catering it to clinical manufacturing processes. With the need for more powerful, precise, and safe genetic editing methods to enhance ACT against solid tumors becoming clear, this BE-mediated dn mutation approach offers a potent new way of armoring CAR T cells against the immunosuppressive TME.

## Materials and Methods

### T cell isolation

Peripheral blood mononuclear cells from de-identified healthy human donors were obtained by automated leukapheresis (StemExpress). CD4+ and CD8+ cells were sorted by immunomagnetic separation using CD4 and CD8 microbeads (Miltenyi Biotec) in combination with a CliniMACS Prodigy cell sorter (Miltenyi Biotec). Signed informed consent was obtained from all donors, and the study was approved by the University of Minnesota Institutional Review Board. All methods were performed in accordance with the relevant guidelines and regulations.

### Cell culture

T cells were cultured in OpTmizer CTS T-Cell basal expansion SFM supplemented with 2.4% OpTmizer CTS T-Cell expansion supplement, 2.3% CTS immune cell serum replacement, L-Glutamine, Penicillin/Streptomycin, 10 mM N-Acetyl-cysteine, 300 IU/mL recombinant IL-2, 5 ng/mL recombinant IL-7, and 5 ng/mL recombinant IL-15 at 37°C and 5% CO_2_. Post-electroporation, T cells were cultured in 24-well G-Rex plates. Raji and OVCAR-8 target cell lines were cultured in RPMI-1640 supplemented with 10% FBS and Penicillin/Streptomycin.

### T cell engineering

Cryopreserved T cells were thawed and stimulated using Dynabeads Human T-Expander CD3/CD28 (ThermoFisher, Waltham, MA) at a 2:1 bead to cell ratio for 48 hours. If applicable, anti-CD19 CAR, anti-Mesothelin CAR, anti-Mesothelin CAR + dnFAS, or anti-Mesothelin CAR + dnTGFβR2 lentivirus was added 24 hours post stimulation at an MOI of 20 with 0.25 mg/mL F108. After the 48 hour stim, Dynabeads were magnetically removed and cells were washed once with PBS prior to resuspension in the appropriate electroporation buffer. For BE-mediated editing, T cells were electroporated with 1 µg of chemically modified sgRNA (IDT, Coralville, Iowa) and 1.5 µg codon optimized ABE8e mRNA (TriLink Biotechnologies, San Diego, CA). T cells were electroporated with the 4D-nucleofector system (Lonza, Basel, Switzerland) using a P3 16-well Nucleocuvette kit, with 1e6 T cells per 20 µL cuvette (program EO-115). T cells were allowed to recover in electroporation cuvettes for 15 minutes before transfer to pre-warmed antibiotic and cytokine-free medium at 37 °C, 5% CO2 for 15 min. Post recovery, T cells were cultured in complete CTS OpTmizer T cell Expansion SFM as described above.

### Sequencing prep and analysis

Seven days after electroporation, genomic DNA was isolated from T cells using Thermo Fisher’s GeneJET genomic DNA purification kit according to the manufacturer’s instructions. Editing efficiency was analyzed by PCR amplification of the targeted loci followed by Sanger sequencing (Eurofins Genomics) (Supplemental Table S1). Sanger sequencing traces of base edited samples were analyzed using EditR (baseeditr.com).

### Fas ligand killing assays

1e6 T cells per condition were exposed to Fas ligand (R&D Systems #126-FL-010) at 300 ng/mL and a conjugating His-Tag antibody (R&D Systems #MAB050) at 10 µg/mL for 24 hours in a 24-well plate. Cell viability was then measured via trypan blue exclusion on a Countess 2 FL cell counting machine at indicated timepoints (ThermoFisher, Waltham, MA).

### TGFβ serial restimulation assays

1e6 T cells per condition were stimulated with plate-bound anti-CD3 antibody (ThermoFisher #16-0037-85) and soluble anti-CD28 antibody (ThermoFisher #16-0289-85) in a 24-well plate for two days. They were then transferred to a clean 24-well plate for two days, and then moved into a 24-well G-Rex plate. After two days in the G-Rex, recombinant TGFβ (Bio-Techne #7754-BH-025) was added at 25 ng/mL. The cells were then allowed to grow for 11 days, media changes performed as needed. TGFβ was replenished every four days for the remainder of the experiment. After the 11 days in the G-Rex, 1e6 per condition were moved to a well of an anti-CD3 antibody coated 24 well plate for another stim with soluble anti-CD28 antibody in the media. After two days, the cells were moved to a 24-well G-Rex plate and allowed to grow for another 11 days, media changes as needed. The process was repeated.

### ICS

T cells were either exposed to Fas ligand (R&D Systems #126-FL-010) at 1.2 µg/mL and a conjugating His-Tag antibody (R&D Systems #MAB050) at 10 µg/mL for 24 hours, or to recombinant TGFβ (Bio-Techne #7754-BH-025) at 100 ng/mL for 30 min. Next, they were exposed to 10 µg/mL brefeldin A (BD Bioscience) and 0.7 µg/mL monensin (BD Bioscience) for 6 hours. Then, they were collected, washed twice with PBS, and incubated with Fixable Viability Dye eFluor780 (ThermoFisher #65-0865-14) and human TruStain FcX (Biolegend #422302) for 30 minutes at 4°C. The cells were then washed with PBS and stained with fluorescently labeled antibodies against CD4, CD8, and CD3 for 30 minutes at 4°C. The cells were then washed again with PBS and permeabilized with Fix/Perm (BD Biosciences) for 20 min at room temperature, washed with Perm/Wash (BD Biosciences), and then stained with fluorescently labeled antibodies against either Smad2/3 (BD Bioscience #562586) or cleaved Caspase 3 (BD Pharmingen #51-68654X) for 30 minutes at 4°C. Post washing, the cells were then analyzed using a CytoFlex S flow cytometer (Beckman Coulter). Data analysis was performed using FlowJo version 10.9.0 (FlowJo LLC).

### Raji serial killing assay

CD19 CAR-T cells were thawed and rested overnight in RPMI-1640 supplemented with 10% FBS and Penicillin/Streptomycin at 37°C and 5% CO2. The next day, recombinant TGFβ (Bio-Techne #7754-BH-025) was added at 25 ng/mL. After another 24 hours, the CAR-T cells were co-cultured with Luciferase-expressing CD19+ Raji target cells at a 3:1 effector:target (E:T) ratio. This was done with 100K target Raji cells per well of a 96-well round-bottom black opaque plate. Each day, D-Luciferin (Goldbio) was added to the media to measure remaining viable Raji target cells via bioluminescence on a plate reader, and afterwards a half media change was performed. Every two days, the TGFβ was replenished. Specific killing was measured as a percentage of luminescence normalized to the control groups (target cells alone), and max-killing was defined as the luminescence value of target cells that had been treated with 1% Triton-X-100. Once maximum killing was reached, the CAR-T were collected and replated with fresh target Raji cells, again at an E:T ratio of 3:1. The reading was then repeated. Each replating, an aliquot of CAR-T was taken for an exhaustion panel via flow cytometry.

### OVCAR-8 killing assays

Mesothelin CAR-T cells engineered with either BE or lentivirus to express dn receptors were thawed and rested overnight in RPMI-1640 supplemented with 10% FBS and Penicillin/Streptomycin. If applicable, recombinant TGFβ (Bio-Techne #7754-BH-025) was added at 25 ng/mL. Luciferase-expressing Mesothelin+ OVCAR-8 target cells were plated at 25K cells per well of an opaque flat-bottom 96-well plate. After 24 hours, the CAR-T cells were co-cultured with the target cells at a 2:1 E:T ratio. Each day for three days, D-Luciferin was added to the media to measure remaining viable Raji target cells via bioluminescence on a plate reader, and afterwards a half media change was performed. On day two, the TGFβ was replenished if applicable. Specific killing was measured as a percentage of luminescence normalized to the control groups (target cells alone), and max-killing was defined as the luminescence value of target cells that had been treated with 1% Triton-X-100.

### Flow cytometry

Approximately 50K - 200K T cells were washed with PBS and then stained in 50 µL PBS with Fixable Viability Dye eFluor780 (ThermoFisher #65-0865-14) and fluorophore-conjugated anti-human antibodies for CD3 (Biolegend #344804), CD4 (Biolegend #300556), CD8 (Biolegend #301042), beta-2-microglobulin (BioLegend #316318), LAG-3 (Biolegend #369212), TIM-3 (Biolegend #345014), and/or PD-1 (Biolegend #329920). The cells were then run on a Beckman Coulter Cytoflex S flow cytometer using BD software, and the data was analyzed on FlowJo version 10.9.0 (FlowJo LLC).

## Data Availability Statement

Datasets available on request from the authors.

## Acknowledgements

B.S.M. is supported by the following NIH grants (R01AI146009, R01AI161017, P01CA254849, P50CA136393, U24OD026641, U54CA232561, P30CA077598, U54CA268069, HT9425-24-1-1005, HT9425-24-1-1002, HT94252410231), and received support from Children’s Cancer Research Fund, the Fanconi Anemia Research Fund, and the Randy Shaver Cancer Research and Community Fund. B.R.W. is supported by the following NIH grants (R21CA237789, R21AI163731, P01CA254849, P50CA136393, U54CA268069, R01AI146009), and received support from Alex’s Lemonade Stand Foundation, Children’s Cancer Research Fund, and Rein in Sarcoma. J.G.S received support from the T32HL007062-46 Hematology Research Training Program and a MIB Agents Young Investigator Award. M.G.K. receives support from F30CA305905-01.

## Author Contributions

Conceptualization: M.G.K., B.R.W. and B.S.M.; Data curation: B.J.W.; Formal analysis, B.J.W.; Funding acquisition: B.R.W. and B.S.M.; Investigation: B.J.W. and E.M.N.; Methodology: B.J.W., N.J.S. and J.G.S.; Project administration: B.R.W. and B.S.M.; Resources: B.R.W. and B.S.M.; Supervision: B.R.W. and B.S.M.; Validation: B.J.W.; Visualization: B.J.W.; Writing – original draft: B.J.W.; Writing – review & editing: B.J.W., M.G.K., N.J.S., J.G.S., E.M.N., B.R.W. and B.S.M. Declarations of Interest: The authors declare no competing interests.

## References

1. Park, J.H., Rivière, I., Gonen, M., Wang, X., Sénéchal, B., Curran, K.J., Sauter, C., Wang, Y., Santomasso, B., Mead, E., et al. (2018). Long-term follow-up of CD19 CAR therapy in acute lymphoblastic leukemia. N. Engl. J. Med. 378, 449–459.

2. Turtle, C.J., Hanafi, L.-A., Berger, C., Hudecek, M., Pender, B., Robinson, E., Hawkins, R., Chaney, C., Cherian, S., Chen, X., et al. (2016). Immunotherapy of non-Hodgkin’s lymphoma with a defined ratio of CD8+ and CD4+ CD19-specific chimeric antigen receptor-modified T cells. Sci. Transl. Med. 8, 355ra116.

3. Porter, D.L., Hwang, W.-T., Frey, N.V., Lacey, S.F., Shaw, P.A., Loren, A.W., Bagg, A., Marcucci, K.T., Shen, A., Gonzalez, V., et al. (2015). Chimeric antigen receptor T cells persist and induce sustained remissions in relapsed refractory chronic lymphocytic leukemia. Sci. Transl. Med. 7, 303ra139.

4. Louis, C.U., Savoldo, B., Dotti, G., Pule, M., Yvon, E., Myers, G.D., Rossig, C., Russell, H.V., Diouf, O., Liu, E., et al. (2011). Antitumor activity and long-term fate of chimeric antigen receptor-positive T cells in patients with neuroblastoma. Blood 118, 6050–6056.

5. Ahmed, N., Brawley, V.S., Hegde, M., Robertson, C., Ghazi, A., Gerken, C., Liu, E., Dakhova, O., Ashoori, A., Corder, A., et al. (2015). Human Epidermal Growth Factor Receptor 2 (HER2) - Specific Chimeric Antigen Receptor-Modified T Cells for the Immunotherapy of HER2-Positive Sarcoma. J. Clin. Oncol. 33, 1688–1696.

6. Lu, Y.-C., Parker, L.L., Lu, T., Zheng, Z., Toomey, M.A., White, D.E., Yao, X., Li, Y.F., Robbins, P.F., Feldman, S.A., et al. (2017). Treatment of Patients With Metastatic Cancer Using a Major Histocompatibility Complex Class II-Restricted T-Cell Receptor Targeting the Cancer Germline Antigen MAGE-A3. J. Clin. Oncol. 35, 3322–3329.

7. Joyce, J.A., and Fearon, D.T. (2015). T cell exclusion, immune privilege, and the tumor microenvironment. Science 348, 74– 80.

8. Rabinovich, G.A., Gabrilovich, D., and Sotomayor, E.M. (2007). Immunosuppressive strategies that are mediated by tumor cells. Annu. Rev. Immunol. 25, 267–296.

9. Wikström, P., Damber, J., and Bergh, A. (2001). Role of transforming growth factor-beta1 in prostate cancer. Microsc. Res. Tech. 52, 411–419.

10. Steiner, M.S., and Barrack, E.R. (1992). Transforming growth factor-beta 1 overproduction in prostate cancer: effects on growth in vivo and in vitro. Mol. Endocrinol. 6, 15–25.

11. Wikström, P., Stattin, P., Franck-Lissbrant, I., Damber, J.E., and Bergh, A. (1998). Transforming growth factor beta1 is associated with angiogenesis, metastasis, and poor clinical outcome in prostate cancer. Prostate 37, 19–29.

12. Abou Shousha, S., Baheeg, S., Ghoneim, H., Zoheir, M., Hemida, M., and Shahine, Y. (2023). The effect of Fas/FasL pathway blocking on apoptosis and stemness within breast cancer tumor microenvironment (preclinical study). Breast Dis. 42, 163–176.

13. Yoshimura, A., and Muto, G. (2011). TGF-β function in immune suppression. Curr. Top. Microbiol. Immunol. 350, 127– 147.

14. Massagué, J. (2008). TGFbeta in Cancer. Cell 134, 215–230.

15. Yamamoto, T.N., Lee, P.-H., Vodnala, S.K., Gurusamy, D., Kishton, R.J., Yu, Z., Eidizadeh, A., Eil, R., Fioravanti, J., Gattinoni, L., et al. (2019). T cells genetically engineered to overcome death signaling enhance adoptive cancer immunotherapy. J. Clin. Invest. 129, 1551–1565.

16. Kloss, C.C., Lee, J., Zhang, A., Chen, F., Melenhorst, J.J., Lacey, S.F., Maus, M.V., Fraietta, J.A., Zhao, Y., and June, C.H. (2018). Dominant-Negative TGF-β Receptor Enhances PSMA-Targeted Human CAR T Cell Proliferation And Augments Prostate Cancer Eradication. Mol. Ther. 26, 1855–1866.

17. Narayan, V., Barber-Rotenberg, J.S., Jung, I.-Y., Lacey, S.F., Rech, A.J., Davis, M.M., Hwang, W.-T., Lal, P., Carpenter, E.L., Maude, S.L., et al. (2022). PSMA-targeting TGFβ-insensitive armored CAR T cells in metastatic castration-resistant prostate cancer: a phase 1 trial. Nat. Med. 28, 724–734.

18. Li, M.O., Sanjabi, S., and Flavell, R.A. (2006). Transforming growth factor-beta controls development, homeostasis, and tolerance of T cells by regulatory T cell-dependent and -independent mechanisms. Immunity 25, 455–471.

19. Gorelik, L., and Flavell, R.A. (2000). Abrogation of TGFbeta signaling in T cells leads to spontaneous T cell differentiation and autoimmune disease. Immunity 12, 171–181.

20. Sweeney, N.P., and Vink, C.A. (2021). The impact of lentiviral vector genome size and producer cell genomic to gag-pol mRNA ratios on packaging efficiency and titre. Mol Ther Methods Clin Dev 21, 574–584.

21. Walther, W., and Stein, U. (2000). Viral vectors for gene transfer: a review of their use in the treatment of human diseases. Drugs 60, 249–271.

22. Gaudelli, N.M., Komor, A.C., Rees, H.A., Packer, M.S., Badran, A.H., Bryson, D.I., and Liu, D.R. (2017). Programmable base editing of A•T to G•C in genomic DNA without DNA cleavage. Nature 551, 464–471.

23. Webber, B.R., Lonetree, C.-L., Kluesner, M.G., Johnson, M.J., Pomeroy, E.J., Diers, M.D., Lahr, W.S., Draper, G.M., Slipek, N.J., Smeester, B.A., et al. (2019). Highly efficient multiplex human T cell engineering without double-strand breaks using Cas9 base editors. Nat. Commun. 10, 5222.

24. Richter, M.F., Zhao, K.T., Eton, E., Lapinaite, A., Newby, G.A., Thuronyi, B.W., Wilson, C., Koblan, L.W., Zeng, J., Bauer, D.E., et al. (2020). Phage-assisted evolution of an adenine base editor with improved Cas domain compatibility and activity. Nat. Biotechnol. 38, 883–891.

25. Strasser, A., Jost, P.J., and Nagata, S. (2009). The many roles of FAS receptor signaling in the immune system. Immunity 30, 180–192.

26. Liu, X., Zhang, Y., Cheng, C., Cheng, A.W., Zhang, X., Li, N., Xia, C., Wei, X., Liu, X., and Wang, H. (2017). CRISPR-Cas9-mediated multiplex gene editing in CAR-T cells. Cell Res. 27, 154–157.

27. Ishigame, H., Mosaheb, M.M., Sanjabi, S., and Flavell, R.A. (2013). Truncated form of TGF-βRII, but not its absence, induces memory CD8+ T cell expansion and lymphoproliferative disorder in mice. J. Immunol. 190, 6340–6350.

28. Lücke, C.D., Philpott, A., Metcalfe, J.C., Thompson, A.M., Hughes-Davies, L., Kemp, P.R., and Hesketh, R. (2001). Inhibiting mutations in the transforming growth factor beta type 2 receptor in recurrent human breast cancer. Cancer Res. 61, 482–485.

29. Yang, J.H., Ki, C.-S., Han, H., Song, B.G., Jang, S.Y., Chung, T.-Y., Sung, K., Lee, H.J., and Kim, D.-K. (2012). Clinical features and genetic analysis of Korean patients with Loeys-Dietz syndrome. J. Hum. Genet. 57, 52–56.

30. Kirmani, S., Tebben, P.J., Lteif, A.N., Gordon, D., Clarke, B.L., Hefferan, T.E., Yaszemski, M.J., McGrann, P.S., Lindor, N.M., and Ellison, J.W. (2010). Germline TGF-beta receptor mutations and skeletal fragility: a report on two patients with Loeys-Dietz syndrome. Am. J. Med. Genet. A 152A, 1016–1019.

31. Loeys, B.L., Chen, J., Neptune, E.R., Judge, D.P., Podowski, M., Holm, T., Meyers, J., Leitch, C.C., Katsanis, N., Sharifi, N., et al. (2005). A syndrome of altered cardiovascular, craniofacial, neurocognitive and skeletal development caused by mutations in TGFBR1 or TGFBR2. Nat. Genet. 37, 275–281.

32. Chung, B.H.-Y., Lam, S.T.-S., Tong, T.M.-F., Li, S.Y.-H., Lun, K.-S., Chan, D.H.-C., Fok, S.F.-S., Or, J.S.-F., Smith, D.K., Yang, W., et al. (2009). Identification of novel FBN1 and TGFBR2 mutations in 65 probands with Marfan syndrome or Marfan-like phenotypes. Am. J. Med. Genet. A 149A, 1452–1459.

33. Kluesner, M.G., Lahr, W.S., Lonetree, C.-L., Smeester, B.A., Qiu, X., Slipek, N.J., Claudio Vázquez, P.N., Pitzen, S.P., Pomeroy, E.J., Vignes, M.J., et al. (2021). CRISPR-Cas9 cytidine and adenosine base editing of splice-sites mediates highly-efficient disruption of proteins in primary and immortalized cells. Nat. Commun. 12, 2437.

34. Wang, M., Krueger, J.B., Gilkey, A.K., Stelljes, E.M., Kluesner, M.G., Pomeroy, E.J., Skeate, J.G., Slipek, N.J., Lahr, W.S., Claudio Vázquez, P.N., et al. (2025). Precision enhancement of CAR-NK cells through non-viral engineering and highly multiplexed base editing. J. Immunother. Cancer 13. 10.1136/jitc-2024-009560.

35. Li, N., Rodriguez, J.L., Yin, Y., Logun, M.T., Zhang, L., Yu, S., Hicks, K.A., Zhang, J.V., Zhang, L., Xie, C., et al. (2024). Armored bicistronic CAR T cells with dominant-negative TGF-β receptor II to overcome resistance in glioblastoma. Mol. Ther. 32, 3522–3538.

36. Liu, X., Zhang, Y., Li, K., Liu, Y., Xu, J., Ma, J., An, L., Wang, H., and Chu, X. (2021). A novel dominant-negative PD-1 armored anti-CD19 CAR T cell is safe and effective against refractory/relapsed B cell lymphoma. Transl. Oncol. 14, 101085.

37. Yi, F., Cohen, T., Zimmerman, N., Dündar, F., Zumbo, P., Eltilib, R., Brophy, E.J., Arkin, H., Feucht, J., Gormally, M.V., et al. (2024). CAR-engineered lymphocyte persistence is governed by a FAS ligand/FAS auto-regulatory circuit. bioRxivorg. 10.1101/2024.02.26.582108.

38. Siegel, P.M., Shu, W., Cardiff, R.D., Muller, W.J., and Massagué, J. (2003). Transforming growth factor beta signaling impairs Neu-induced mammary tumorigenesis while promoting pulmonary metastasis. Proc. Natl. Acad. Sci. U. S. A. 100, 8430–8435.

39. Schmidt, R., Ward, C.C., Dajani, R., Armour-Garb, Z., Ota, M., Allain, V., Hernandez, R., Layeghi, M., Xing, G., Goudy, L., et al. (2024). Base-editing mutagenesis maps alleles to tune human T cell functions. Nature 625, 805–812.

40. Liu, Q., Fang, L., and Wu, C. (2022). Alternative splicing and isoforms: From mechanisms to diseases. Genes (Basel) 13, 401.

