## Supplemental figures and table for "Installation of Dominant-Negative Mutations in *FAS* and *TGFβR2* via Base Editing in Primary T Cells"

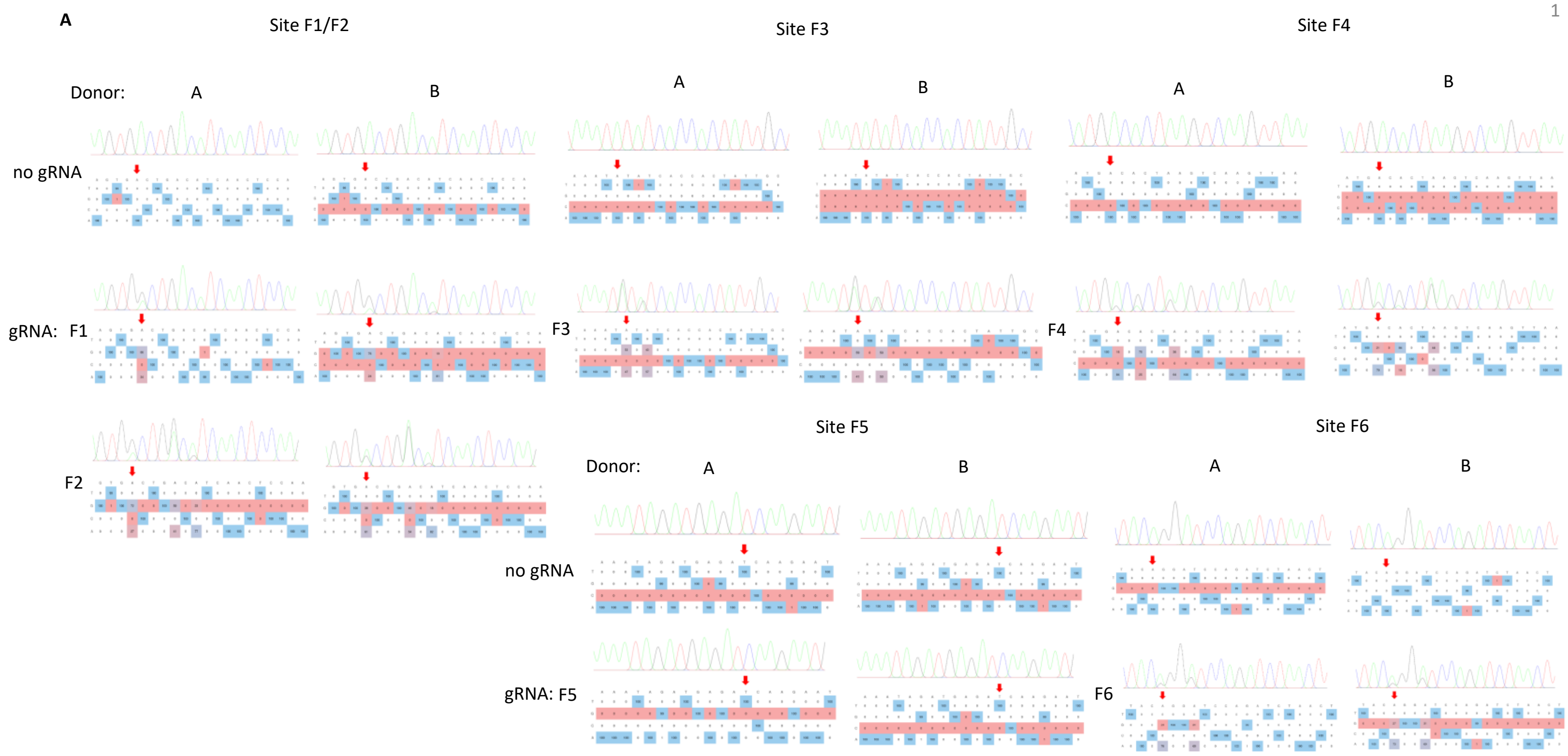


**Figure S1: ABE8e efficiency installs mutations in *FAS*.** (**A**) Representative chromatograms for Sanger sequencing of target loci seven days after guide RNAs were electroporated into T cells with ABE8e mRNA. These chromatograms represent the T cells used in the initial sequencing experiment (Figure 1C).

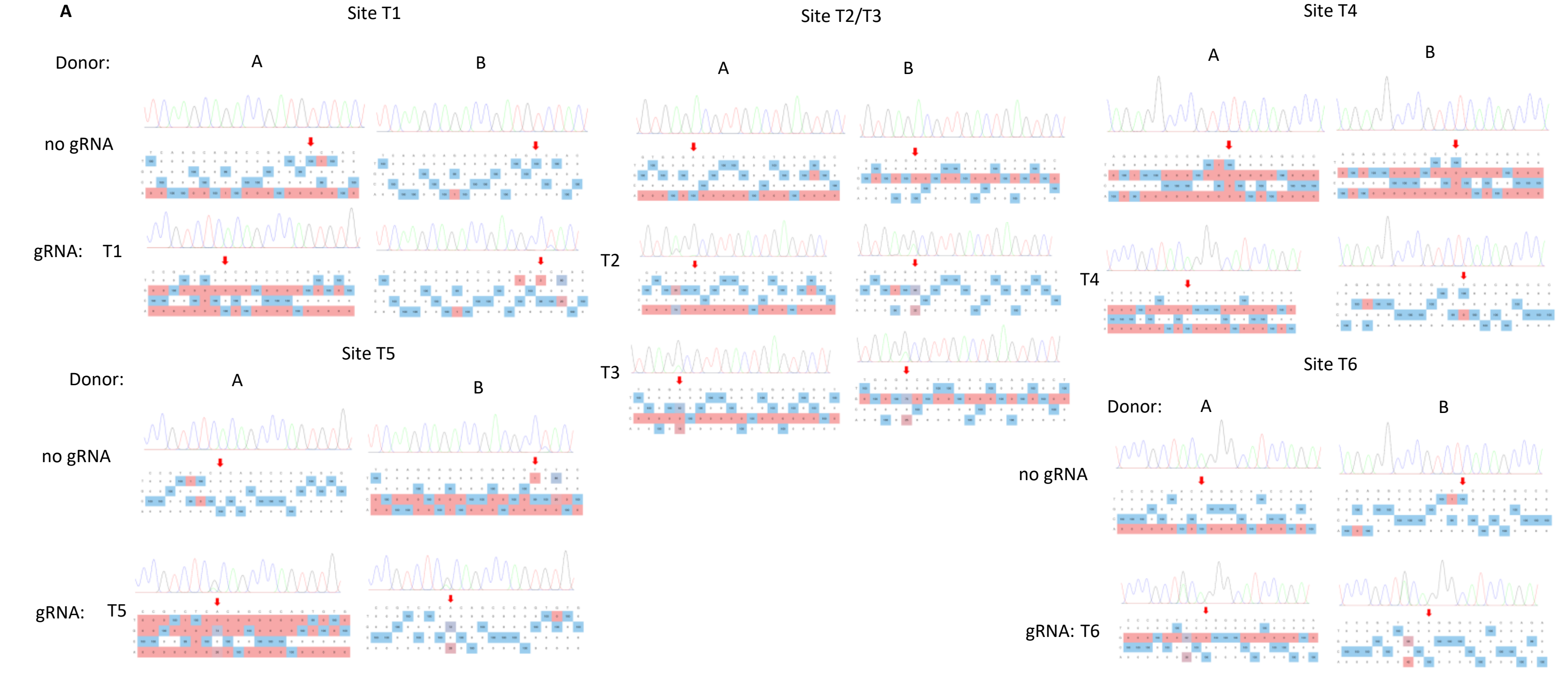


**Figure S2: ABE8e efficiency installs mutations in *TGFβR2*.** (**A**) Representative chromatograms for Sanger sequencing of target loci seven days after guide RNAs were electroporated into T cells with ABE8e mRNA. These chromatograms represent the T cells used in the initial sequencing experiment (Figure 1C).

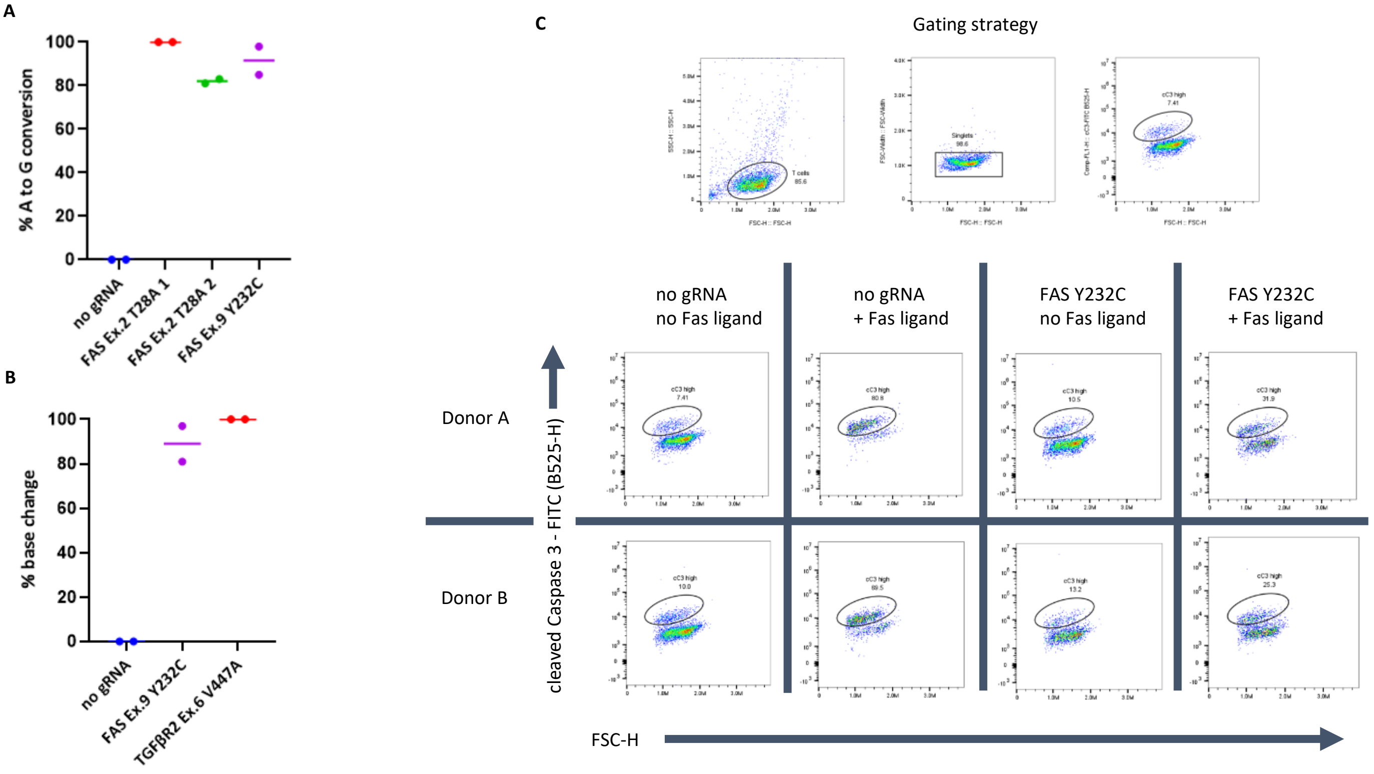


**Figure S3: dnFAS T cells exhibit lower cC3 staining after exposure to FASL.** (**A**) Base editing activity for gRNAs used in the Fas ligand killing assay. (**B**) Base editing activity for gRNAs used in the ICS assays. Guide RNAs were electroporated into T cells with ABE8e mRNA and base editing activity was assessed seven days later via sanger sequencing. (n = 2 donors). (**C**) Representative flow plots for Fas ICS assay. T cells were either exposed to Fas ligand and a conjugating His-Tag antibody for 24 hours or just the His-Tag antibody and then ICS was performed for cleaved Caspase 3.

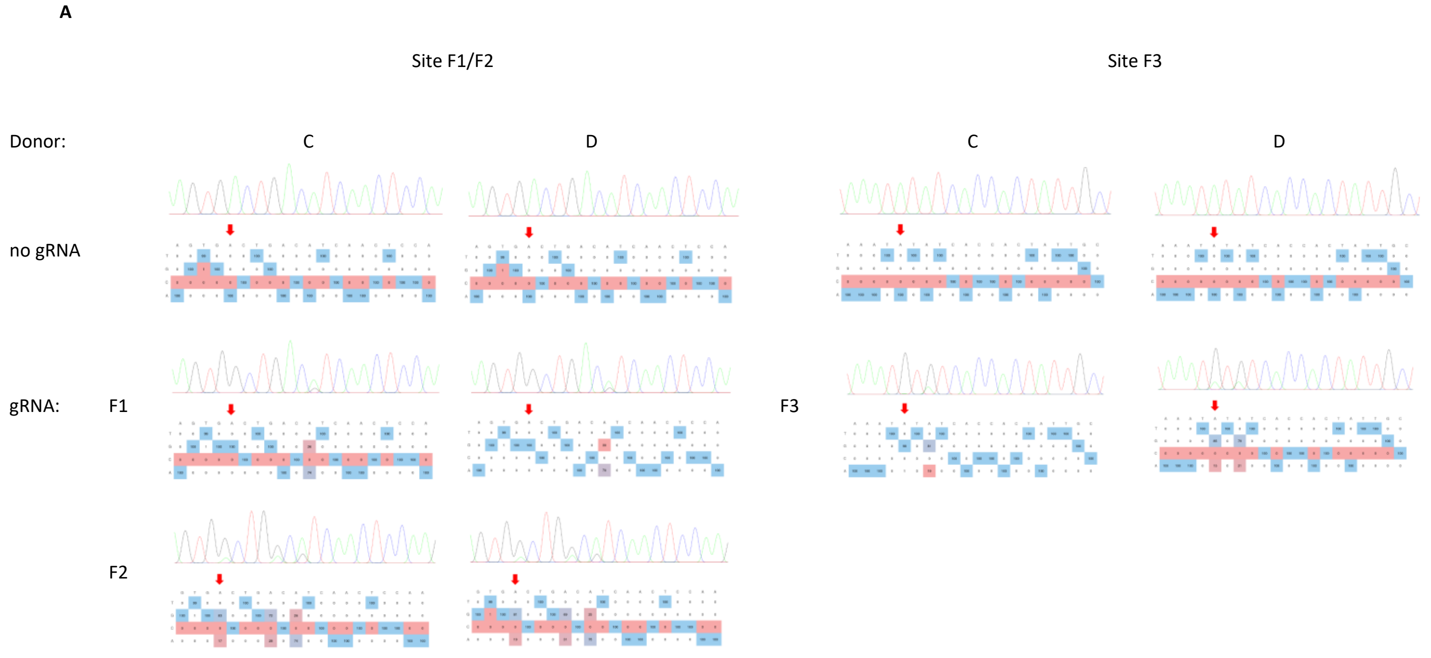


**Figure S4: ABE8e efficiency installs candidate dn mutations in *FAS*.** (**A**) Representative chromatograms for Sanger sequencing of target loci seven days after guide RNAs were electroporated into T cells with ABE8e mRNA. These chromatograms represent the T cells used in the Fas ligand-based killing assay (Supplemental Figure 3A).

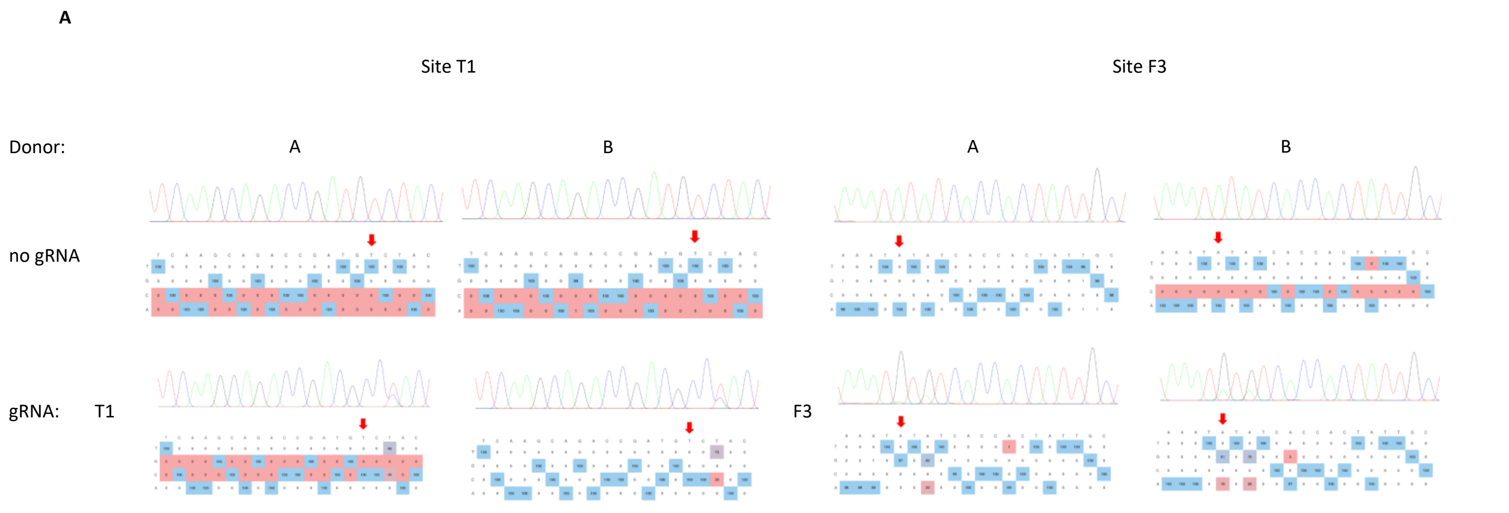


**Figure S5: ABE8e efficiency installs *FAS* Y232C and *TGFβR2* V447A dn mutations.** (**A**) Representative chromatograms for Sanger sequencing of target loci seven days after guide RNAs were electroporated into T cells with ABE8e mRNA. These chromatograms represent the T cells used in the ICS assays (Figure 2C and Figure 3D).

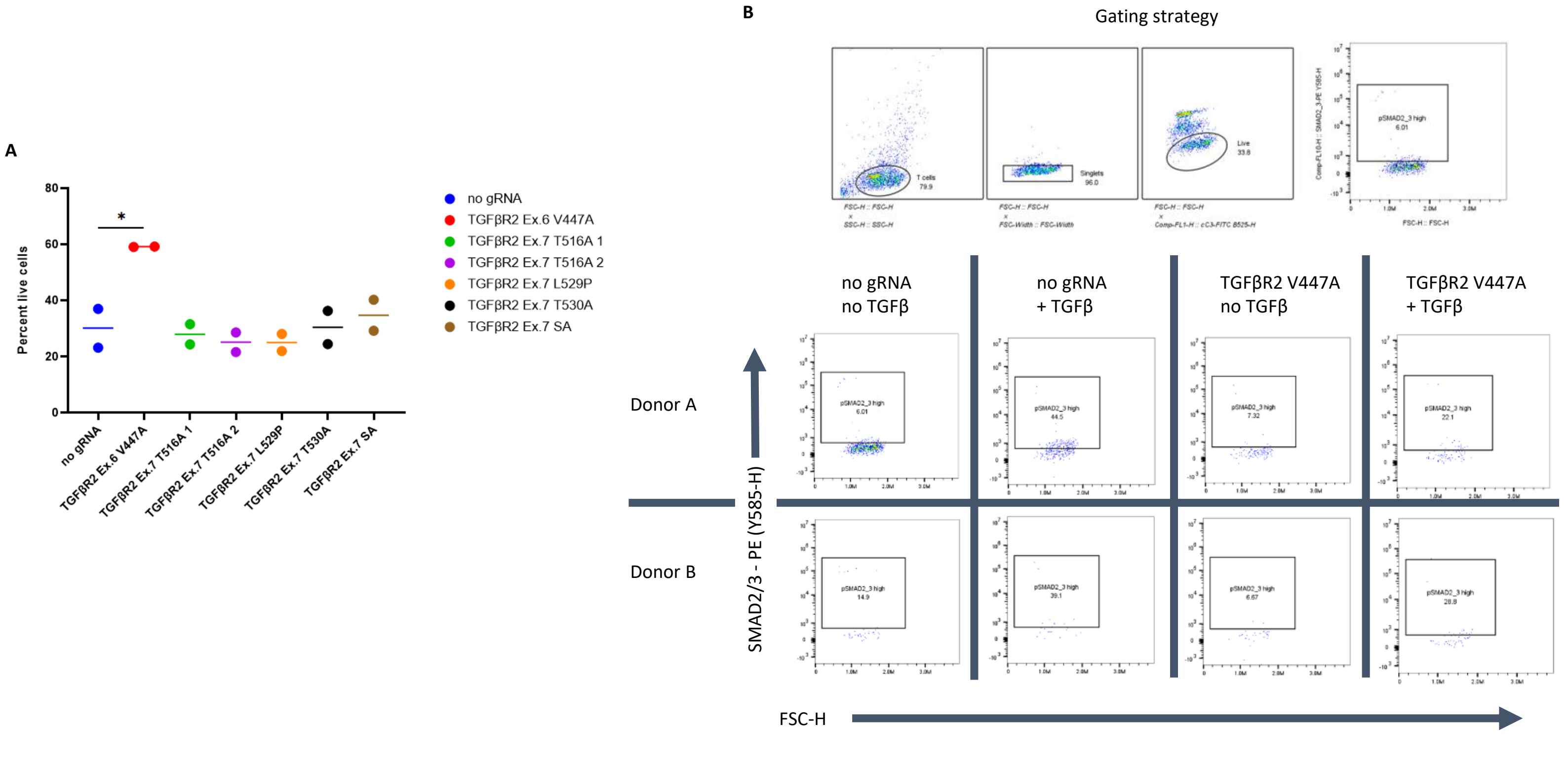


**Figure S6: dnTGFβR2 T cells exhibit lower pSMAD2/3 staining and higher viability after exposure to TGFβ.** (**A**) Percent live cells four days after the third stimulation in the serial plate stim experiment. *(n = 2 donors). ns = not significant, *p<0.05, **p<0.01, ***p<0.001, ****p<0.0001, One-way ANOVA with Dunnett's multiple comparisons test against no gRNA control.*(**B**) Representative flow plots for TGFβ ICS assay. T cells were either exposed to TGFβ for 30 minutes or media alone and then ICS was performed for SMAD2/3

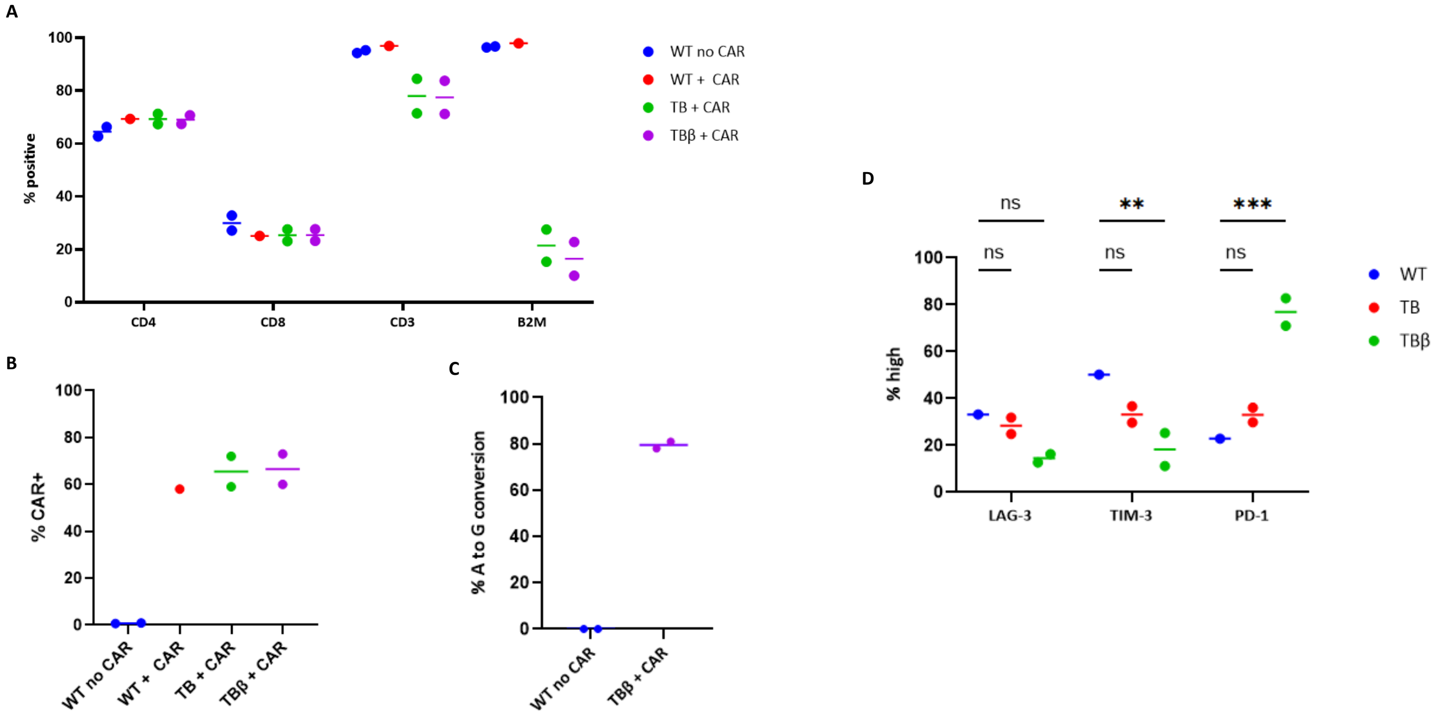


**Figure S7: dnTGFβR2 engineering is easily added to clinically-relevant CAR-T manufacturing processes.** (**A**) Flow cytometry data for assessing knock out efficiencies on T cells used in the TGFβ serial killing assay on day 7 post-engineering. (**B**) Flow cytometry data for assessing CAR integration on T cells used in the TGFβ serial killing assay on day 7 post-engineering. (**C**) Sequencing data for base editing activity on T cells used in the TGFβ serial killing assay on day 7 post-engineering. Desired edit is TGFβR2 V447A. (**D**) Inhibitory receptor panel done on T cells before the second replating (day 9) during the TGFβ serial killing assay. WT = wild type (no knock-outs). TB = TRAC KO and B2M KO. TBβ = TRAC KO, B2M KO, and TGFβR2 V447A. (n = 2 donors). *ns = not significant, *p<0.05, **p<0.01, ***p<0.001, ****p<0.0001, (D) Two-way ANOVA with Uncorrected Fisher's LSD.*

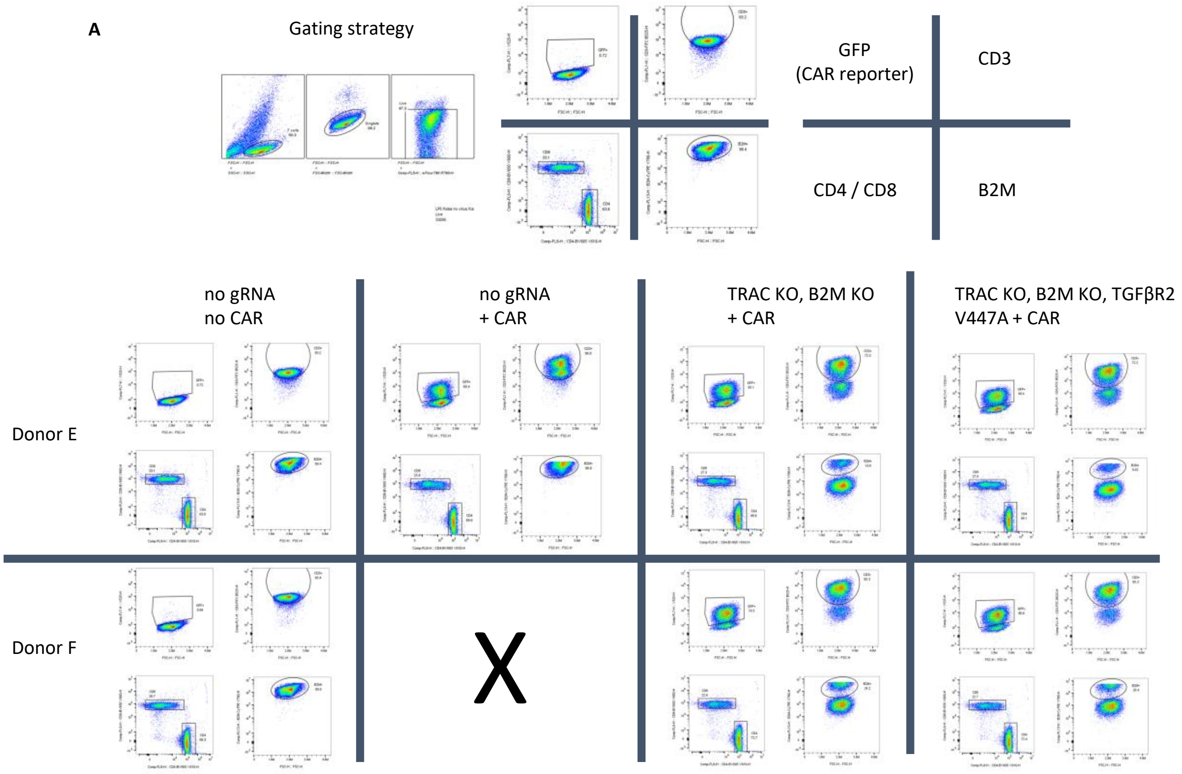


**Figure S8: dnTGFβR2 engineering does not affect other CAR T cell edits.** (**A**) Representative flow plots for TGFβ serial killing assay T cell production. Flow cytometry was done seven days post T cell engineering.

**A**

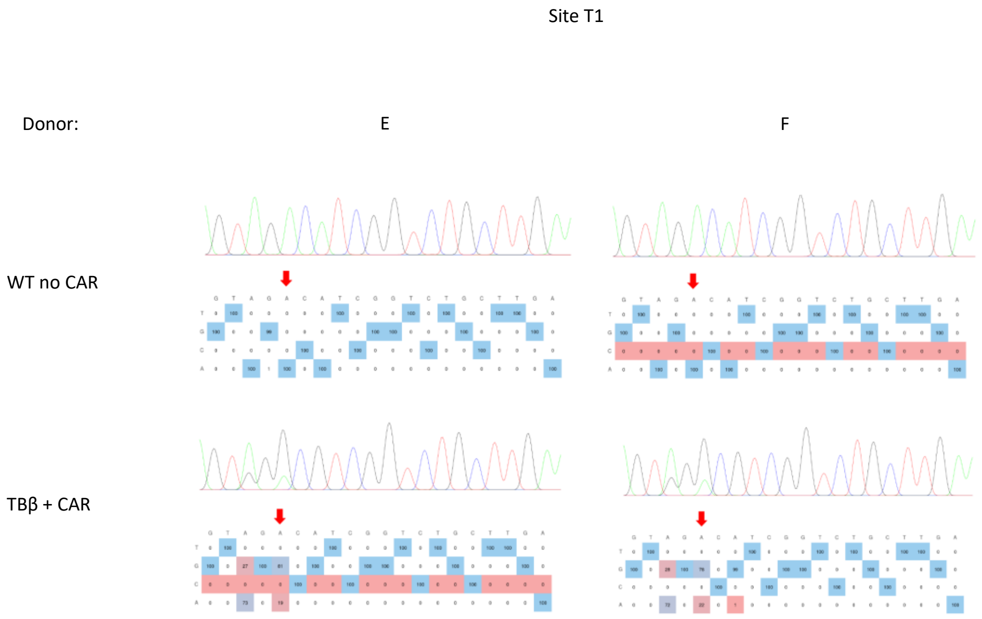


**Figure S9: ABE8e efficiency installs *TGFβR2* V447A alongside *TRAC* and *B2M* KO.** (**A**) Representative chromatograms for Sanger sequencing of target loci seven days after guide RNAs were electroporated into T cells with ABE8e mRNA. These chromatograms represent the T cells used in the TGFβ serial killing assay (Supplemental Figure 7C).

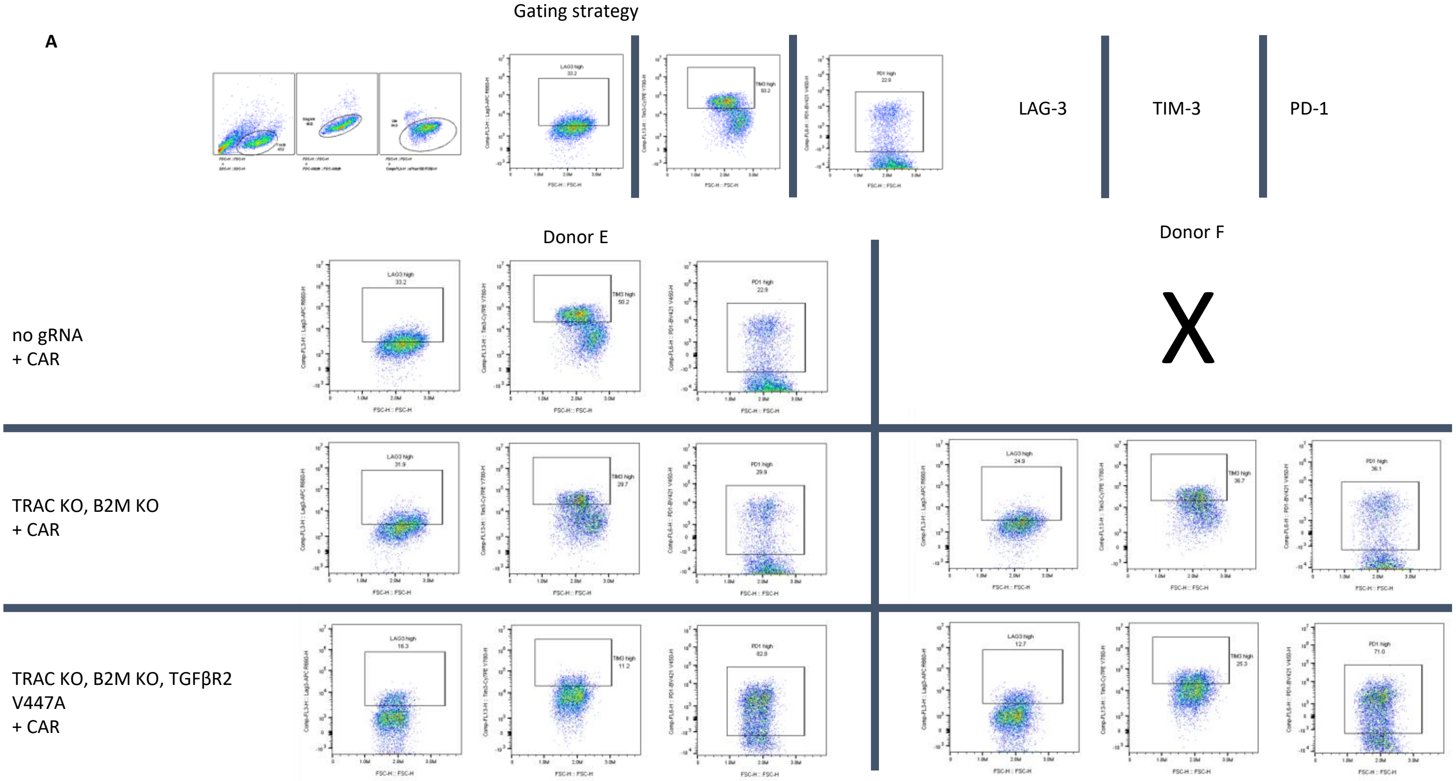


**Figure S10: dnTGFβR2 CAR T cells exhibit less of an exhausted phenotype during serial killing assay.** (**A**) Representative flow plots for TGFβ serial killing assay exhaustion panel. The exhaustion panel was done before the second replating, on day 9 of the serial killing assay.

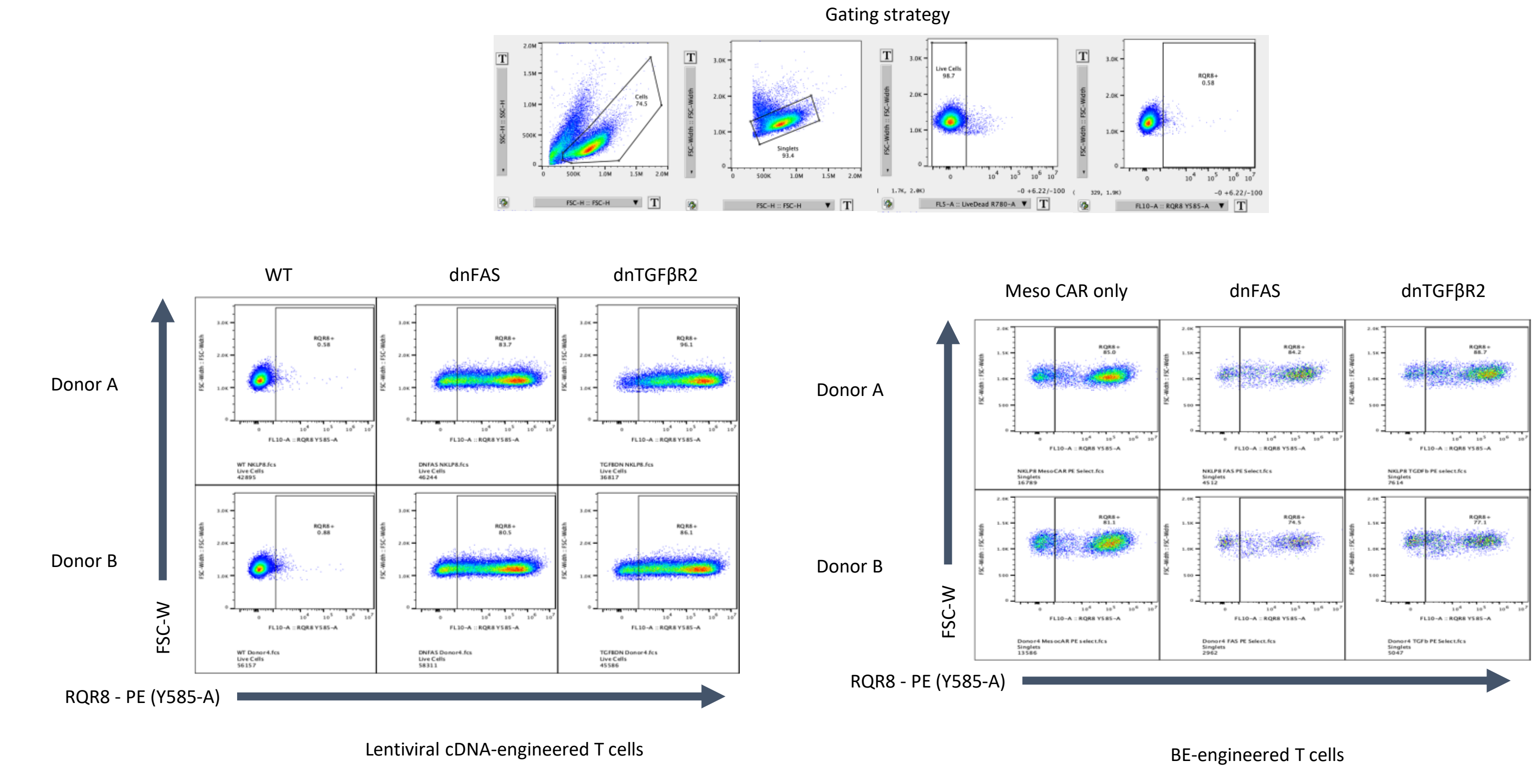


**Figure S11: CAR integration efficiency is not impacted by the simultaneous BE installation of dnFAS and dnTGFβR2 mutations in T cells.** Representative flow plots showing RQR8 reporter expression for lentiviral cDNA and BE-based engineering.

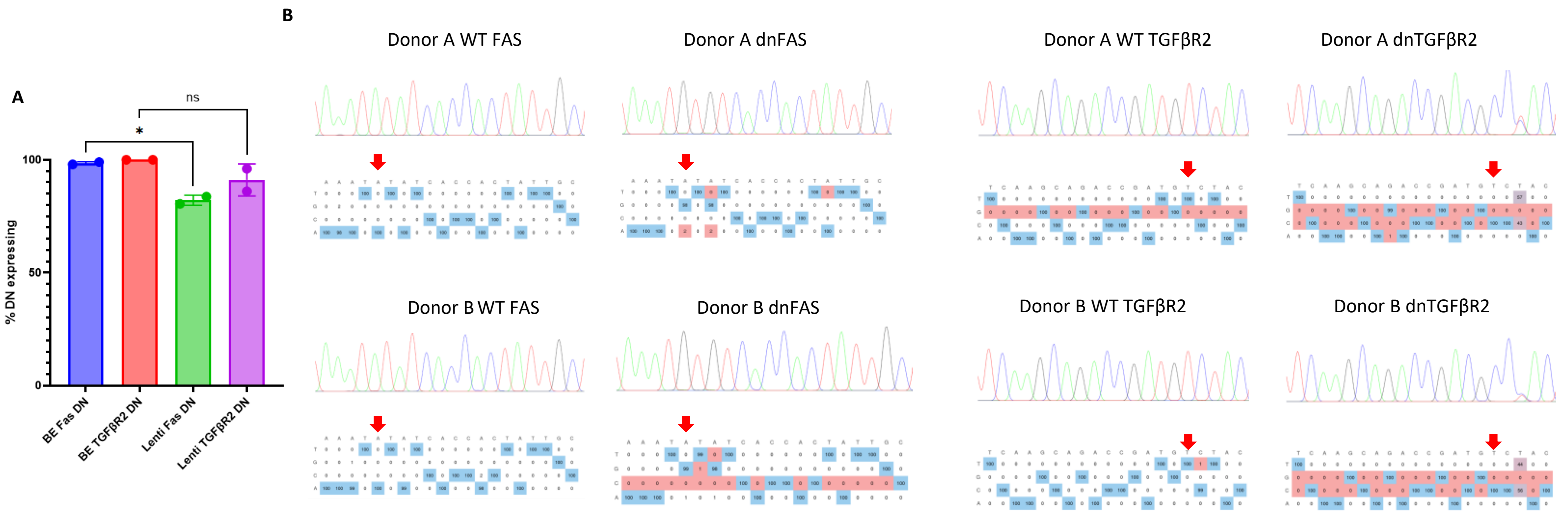


**Figure S12: BE can introduce dnFAS and dnTGFβR2 mutations with equal or higher efficiency than lentiviral cDNA.** (**A**) Percent dn receptor expression for BE and lentiviral cDNA-engineered CAR T cells. (**B**) Representative sanger sequencing chromatograms for BE-engineered CAR T cells.

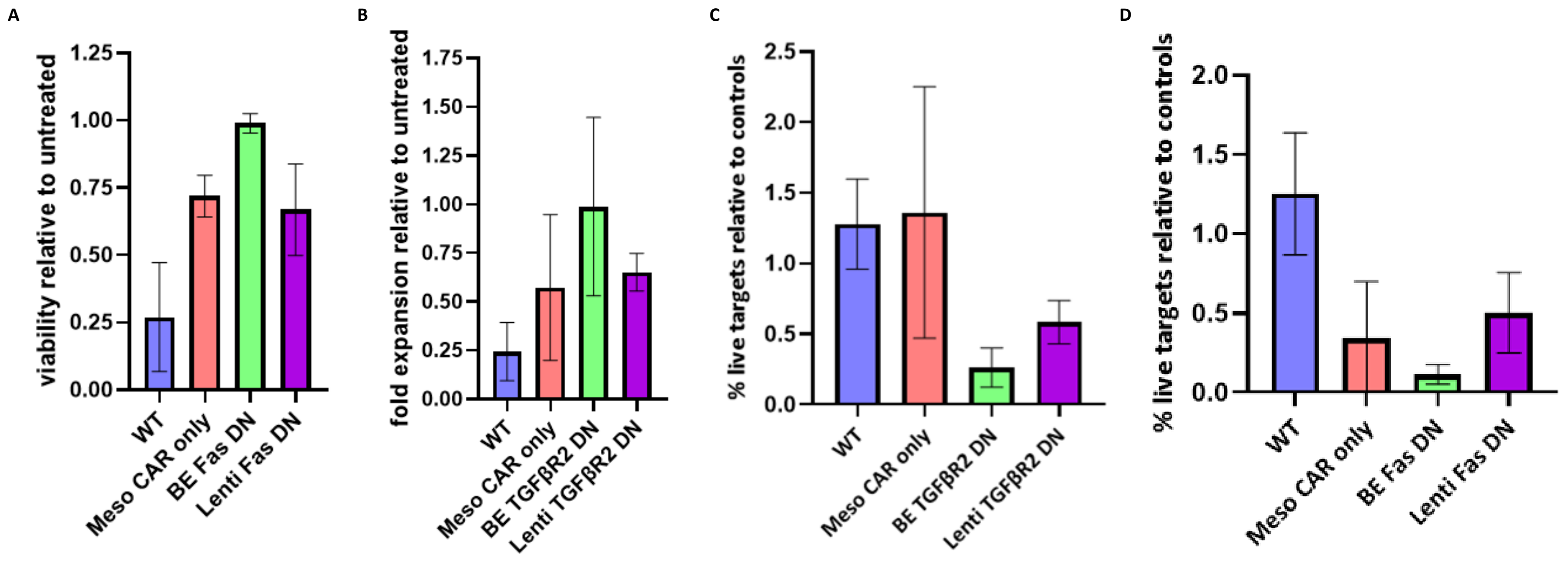


**Figure S13: BE-engineered dnFAS and dnTGFβR2 CAR T cells perform similar or superior to lentiviral cDNA-engineered counterparts in *in vitro* functional assays.** (**A**) Percent viable cells remaining 24 hours after Fas-ligand killing assay. (**B**) Fold expansion after the second stim in a TGFβ serial plate restim. (**C**) Remaining target cells as measured via bioluminescence after 72 hours in a co-culture killing assay with TGFβ supplemented in the media. (**D**) Remaining target cells as measured via bioluminescence after 72 hours in a co-culture killing assay. *(n = 2 donors).*

**Table S1: gRNA and primer sequences.**

| Shorthand name | Full name | Sequence (5' to 3') | Forward sequencing primer sequence (5' to 3') | Reverse sequencing primer sequence (5' to 3') | Primer set |
|---|---|---|---|---|---|
| F1 | FAS Ex.2 T28A_1 | AGTGACTGACATCAACTCCA | GTGGAGCCCTCACATTGTCT | AACCACATCAAATAAGCGTGA | 1 |
| F2 | FAS Ex.2 T28A_2 | GTGACTGACATCAACTCCAA | GTGGAGCCCTCACATTGTCT | AACCACATCAAATAAGCGTGA | 1 |
| F3 | FAS Ex.9 Y232C | AAATATATCACCACTATTGC | TTCCCCTAGTCAGCTCTTCA | CCAAGCTTTGGATTTCATTTC | 2 |
| F4 | FAS Ex.9 T241A | ATGACACTAAGTCAAGTTAA | TTCCCCTAGTCAGCTCTTCA | CCAAGCTTTGGATTTCATTTC | 2 |
| F5 | FAS Ex.9 I262T | ATTCTTGATCTCATCTATTT | TTCCCCTAGTCAGCTCTTCA | CCAAGCTTTGGATTTCATTTC | 2 |
| F6 | FAS Ex.6 SA | TACAGGATCCAGATCTAACT | ACACTCACCAGCAACACCAA | ATACCACTCAATGCCCCAAA | 3 |
| T1 | TGFBR2 Ex.6 V447A | GTAGACATCGGTCTGCTTGA | GGGTTTTCAGGGAGAGAACA | ACTGCTTTGTAACCCCTGGA | 4 |
| T2 | TGFBR2 Ex.8 T516A_1 | GTGAGACGTTGACTGAGTGC | CAGGCACTCAGTCAGCACAT | TCTGCTTATCCCCACAGCTT | 5 |
| T3 | TGFBR2 Ex.8 T516A_2 | TGAGACGTTGACTGAGTGCT | CAGGCACTCAGTCAGCACAT | TCTGCTTATCCCCACAGCTT | 5 |
| T4 | TGFBR2 Ex.8 L529P | GGGCTGTGAGACGGGCCTCT | CAGGCACTCAGTCAGCACAT | TCTGCTTATCCCCACAGCTT | 5 |
| T5 | TGFBR2 Ex.8 T530A | CCGTCTCACAGCCCAGTGTG | CAGGCACTCAGTCAGCACAT | TCTGCTTATCCCCACAGCTT | 5 |
| T6 | TGFBR2 Ex.8 SA | CCCGCTACAGGGCATCCAGA | CAGGCACTCAGTCAGCACAT | TCTGCTTATCCCCACAGCTT | 5 |
